# Natural isolate and recombinant SARS-CoV-2 rapidly evolve in vitro to higher infectivity through more efficient binding to heparan sulfate and reduced S1/S2 cleavage

**DOI:** 10.1101/2021.06.28.450274

**Authors:** Nikita Shiliaev, Tetyana Lukash, Oksana Palchevska, David K. Crossman, Todd J. Green, Michael R. Crowley, Elena I. Frolova, Ilya Frolov

**Affiliations:** Department of Microbiology, University of Alabama at Birmingham, Birmingham, Alabama, USA; UAB Genomics Core Facility, Department of Genetics, University of Alabama at Birmingham, Birmingham, Alabama, USA

## Abstract

One of the severe acute respiratory syndrome coronavirus 2 (SARS-CoV-2) virulence factors is the ability to interact with high affinity to the ACE2 receptor, which mediates viral entry into cells. The results of our study demonstrate that within a few passages in cell culture, both the natural isolate of SARS-CoV-2 and the recombinant, cDNA-derived variant acquire an additional ability to bind to heparan sulfate (HS). This promotes a primary attachment of viral particles to cells before their further interactions with the ACE2. Interaction with HS is acquired through multiple mechanisms. These include i) accumulation of point mutations in the N-terminal domain (NTD) of the S protein, which increase the positive charge of the surface of this domain, ii) insertions into NTD of heterologous peptides, containing positively charged amino acids, and iii) mutation of the first amino acid downstream of the furin cleavage site. This last mutation affects S protein processing, transforms the unprocessed furin cleavage site into the heparin-binding peptide and makes viruses less capable of syncytia formation. These viral adaptations result in higher affinity of viral particles to heparin sepharose, dramatic increase in plaque sizes, more efficient viral spread, higher infectious titers and two orders of magnitude lower GE:PFU ratios. The detected adaptations also suggest an active role of NTD in virus attachment and entry. As in the case of other RNA+ viruses, evolution to HS binding may result in virus attenuation *in vivo*.

**IMPORTANCE:** The spike protein of SARS-CoV-2 is a major determinant of viral pathogenesis. It mediates binding to ACE2 receptor and later, fusion of viral envelope and cellular membranes. The results of our study demonstrate that SARS-CoV-2 rapidly evolves during propagation in cultured cells. Its spike protein acquires mutations in the N-terminal domain (NTD) and in P1‘ position of the furin cleavage site (FCS). The amino acid substitutions or insertions of short peptides in NTD are closely located on the protein surface and increase its positive charge. They strongly increase affinity of the virus to heparan sulfate, make it dramatically more infectious for the cultured cells and decrease GE:PFU ratio by orders of magnitude. The S686G mutation also transforms the FCS into the heparin-binding peptide. Thus, the evolved SARS-CoV-2 variants efficiently use glycosaminoglycans on the cell surface for primary attachment before the high affinity interaction of the spikes with the ACE2 receptor.

## INTRODUCTION

Severe acute respiratory syndrome coronavirus 2 (SARS-CoV-2) has recently emerged in Wuhan, China and then demonstrated an unprecedented spread all over the world. To date, it is circulating on all continents, and the associated COVID-19 disease has already caused millions of deaths worldwide (1, 2).

SARS-CoV-2 is a member of the *Betacoronavirus* genus (β-CoV) in the *Coronaviridae* family (3). It is a spherical, lipid envelope-containing virus. SARS-CoV-2 genome (G RNA) is represented by a single-stranded RNA of positive polarity, ∼30 kb in length. It mimics cellular mRNA by having a Cap and a poly(A)-tail at the 5’ and 3’ termini, respectively. Viral nonstructural proteins (nsps) are translated directly from G RNA as two overlapping polyproteins ORF1a and ORF1ab, which are further processed into individual nsp1-to-16 by the encoded viral proteases. These nsps function in replication of the viral genome and synthesis of 8 subgenomic RNAs (SG RNAs), which encode virus-specific structural and accessory proteins (4). Functions of the structural N protein include packaging of viral G RNA during formation of infectious viral particles. It is also required for synthesis of G RNA during viral replication. M and E proteins are imbedded into the viral lipid envelope and determine virion assembly and release. The structural S protein forms trimeric spikes on the surface of virions and is a major determinant of viral pathogenesis. It mediates binding to the ACE2 receptor and, later, fusion of the viral envelope and cellular membranes, resulting in release of G RNA-containing nucleocapsid into the cytoplasm. SARS-CoV-2 utilizes two entry mechanisms. In the first (early) pathway, the transmembrane serine protease(s), such as TMPRSS2, mediates cleavage of the S2’ site, which releases the fusion peptide, and this ultimately leads to fusion of viral envelope and the plasma membrane. In the cells lacking TMPRSS proteases, the S2’ processing is achieved by cathepsins, and fusion takes place in the endosomes (5, 6). The distinguishing feature of the SARS-CoV-2-specific S protein is that it contains an additional furin cleavage site, which mediates processing of this protein into S1/S2 subunits before particle release from infected cells (7).

Human ACE2 is a high affinity receptor of SARS-CoV-2 and determines specificity of the infection. However, similar to numerous other viral infections, interaction with the receptor may not be the only determinant of virus attachment and entry (8). Other attachment factors at the plasma membrane may also play an important role(s). Despite the low affinity of this binding to additional factors at the membrane, the high abundance of such molecules may promote the primary attachment of viral particles before their further interactions with receptors. In many cases, these additional attachment factors have been identified as glycosaminoglycans (GAGs), most frequently heparin and heparan sulfate (HS). They are ubiquitously expressed at the plasma membrane of most of vertebrate cell lines and in the extracellular matrix (9). It appears that SARS-CoV-2 is not an exception. The receptor-binding domain of its S protein was found to interact with HS (10). This binding changes the S protein receptor-binding domain (RBD) from a closed to an open conformation, thereby promoting more efficient binding to ACE2.

The ability of the viruses to interact with HS does not exclude their further evolution in developing a more efficient binding of their structural proteins to HS and other GAGs. For example, serial passaging of alphaviruses and flaviviruses in cultured cells during development of live attenuated vaccines results in accumulation of a very few point mutations, which increase a number positively charged amino acids (aa) on the glycoprotein surface (11–15). These aa substitutions increase affinity of the virus to HS, viral infectivity *in vitro* and, compared to natural isolates, strongly decrease the genome equivalent to plaque forming units (GE:PFU) ratio. On the other hand, these mutations usually result in variants that are dramatically less pathogenic, but the mechanism of attenuation remains incompletely understood. However, in some cases, HS-binding alphaviruses exhibit high neurovirulence in mice (15, 16).

The results of our new study show that within very few passages *in vitro*, SARS-CoV-2 rapidly evolves, and variants demonstrating more efficient spread in cultured cells are selected. These variants acquire higher affinities to HS, which largely result from the insertion of a short positively charged peptide into the N-terminal domain (NTD) of the S protein. Another means of adaptation to cell culture is inactivation of SARS-CoV-2-specific furin cleavage site by substitution of a single aa. This mutation appears rapidly during virus rescue from infectious cDNA in Vero E6 cells and becomes prevalent. The positively charged amino acids in the uncleaved furin site also increase the affinity of the virus to HS. However, the decrease in cleavage efficiency does not prevent further evolution that results in accumulation of additional positively charged amino acids in the NTD. Both, the insertion of the peptide and point mutation in the furin cleavage site, strongly increase infectious titers and plaque size of SARS-CoV-2 on Vero cells and accelerate viral spread. They also decrease GE:PFU ratio by several orders of magnitude.

## MATERIALS AND METHODS

### Cell cultures

The BHK-21 cells were kindly provided by Paul Olivo (Washington University, St. Louis, MO). The NIH 3T3, Calu-3 and Vero E6 cells were obtained from the American Type Culture Collection (Manassas, VA). Huh7.5 cells were kindly provided by Charles Rice (Rockefeller University, New York, NY). BHK-21, Vero E6, NIH 3T3 cells and their derivatives were maintained in alpha minimum essential medium supplemented with 10% fetal bovine serum (FBS) and vitamins. Huh7.5 and Calu-3 cells were maintained in Dulbecco’s modified Eagle medium supplemented with 10% (FBS).

### SARS-CoV2 isolates

The stock of SARS-CoV-2 strain 2019-nCoV/USA_WA1/2020 was derived from the first patient diagnosed in the US. This virus isolate was originally provided by Dr. Natalie Thornburg from the Centers for Disease Control and Prevention in Atlanta, GA (17), and amplified on Vero E6 cells at the World Reference Center for Emerging Viruses and Arboviruses (WRCEVA) at the University of Texas Medical Branch at Galveston (UTMB). Passage 4 was received from WRCEVA and further passaged in this study. In the following sections this isolate is referred to as WA1/2020. The second isolate (CoV2/UAB) was kindly provided by Dr. William Britt (The University of Alabama at Birmingham, USA), and passage 1 was used in this study. The S protein of this isolate contained a single mutation D614G, which is relatively common for SARS-CoV-2 isolates (18).

### Plasmid constructs

The cDNA of SARS-CoV-2 (NC_045512.2) genome was assembled in two fragments (5’ and 3’ fragments), which were cloned into low copy number plasmids pACNR1180 (19) and Proteus1 (20), and propagated in *E.coli* WM1100. Fragments were assembled from gene blocks synthesized by Integrated DNA Technologies (IDT). The 7a gene in the 3’ fragment was replaced by the GFP-coding sequence. The 5’ fragment was positioned under the control of the SP6 promoter.

Sequences of both cDNA fragments were verified by Sanger sequencing. Mutations were introduced into the S protein-coding sequence by cloning corresponding gene blocks to replace the original fragments. Human ACE2 gene (hACE2) was kindly provided by Luis Martinez-Sobrido (Texas Biomedical Research Institute, San Antonio, TX). It was cloned into a modified PiggyBac plasmid under control of the CMV promoter (PiggyBAC/ACE2). The N protein-coding sequence was cloned into a noncytopathic replicon of Venezuelan equine encephalitis virus under the control of the subgenomic promoter (VEErep/N/Pac) (21). The second promoter was driving expression of the selectable marker, puromycin acetyl transferase (Pac).

### Stable cell lines

Stable ACE2-expressing cell lines were generated by co-transfecting BHK-21, NIH 3T3, Vero E6 and Huh7.5 cells with PiggyBAC/ACE2 and integrase-encoding plasmids (System Biosciences, Inc.) using Lipofectamine 3000, according to the protocol recommended by the manufacturer (Invitrogen). Blasticidin selection was used to select stable ACE2-expressing cell lines. A stable, N protein-expressing cell line was generated by electroporation of the *in vitro*-synthesized VEErep/N/Pac RNA into BHK-21 or into BHK/ACE2 cells. Electroporated cells were further passaged in the presence of 5 μg/ml of puromycin in the media.

**Infectious titers** of SARS-CoV-2 were determined by plaque assay on Vero E6 or Vero/ACE2 cells. Cells were seeded in 6-well Costar plates (2.5×10^5^ cells/well). In 4 h, they were infected by serial dilutions of the viruses. After 30 min of incubation at 37°C, cells were covered with 2 ml of 0.5% agarose supplemented with DMEM and 3% FBS. After 60-70 h of incubation at 37°C, cells were fixed by 10% formaldehyde and then stained with crystal violet.

### Rescuing of the recombinant viruses

Plasmids encoding cDNAs of viral genomes were purified by ultracentrifugation in CsCl gradients. The 5’ fragment-containing plasmid was digested by SacII, located in the plasmid backbone, dephosphorylated by alkaline phosphatase and after its inactivation, digested by SacI. The fragment encoding the SP6 promoter and ∼16,000 nt of the 5’ end of viral genome was isolated by agarose gel electrophoresis. Plasmid containing the 3’ fragment of the genome was digested by SacI and SfiI, whose site is located downstream of the poly(A) tail, and the 3’ fragment of the viral sequence was also isolated by agarose gel electrophoresis. The 5’ and 3’ fragments (2 μg of each) were ligated for 1 h at 16°C by T4 DNA ligase (Invitrogen) and purified by phenol-chloroform extraction. RNAs were synthesized *in vitro* by SP6 RNA polymerase (New England Biolabs) in the presence of a cap analog (New England Biolabs) under the conditions recommended by the manufacturer. The qualities of the RNAs were evaluated by electrophoresis in nondenaturing agarose gels. RNAs were used for electroporation of cells without additional purification.

The *in vitro*-synthesized RNAs (∼1 μg) were electroporated into BHK-21 or BHK/ACE2 cells containing VEErep/N/Pac under previously described conditions (22, 23). Electroporated cells were seeded onto subconfluent monolayers of Vero/ACE2 cells and incubated at 37°C. Depending on the construct, viruses were harvested at 48 or 72 h post electroporation, and titers were determined by a plaque assay on Vero/ACE2 cells.

### Analysis of virus binding to heparin sepharose

For this analysis, virus stocks were prepared in serum-free VP-SF media (Invitrogen). Viruses were harvested before CPE development, additionally clarified by centrifugation at 16,000 x g, and supernatants were used for further analysis. Column of heparin sepharose (0.4 ml) was equilibrated with 0.66 x PBS (NaCl concentration of 0.1 M). Viral samples (1 ml) were diluted by 0.5 volume of water to reduce concentration of NaCl to 0.1 M and loaded on to the column in the biosafety cabinet. Then column was washed by a step gradient of PBS containing NaCl at concentrations increasing from 0.1 to 0.5 M, and viral titers were determined in each fraction, including flow through (FT), by plaque assay on Vero/ACE2 cells.

### Analysis of spike protein processing in the released virions

Viral samples were prepared in serum-free VP-SF media. To avoid contamination with cell debris, they were harvested before CPE development. Virus was inactivated by adding 4x protein gel loading buffer and analyzed by Western blotting using S2- and N protein-specific Abs (Sinobiological and ProSci, respectively) and corresponding secondary Abs labeled with infrared dyes. The membranes were scanned on an Odyssey Imaging System (LI-COR Biosciences).

### Next generation sequencing (NGS)

G RNAs were isolated from the viral samples using Direct-zol™ RNA Miniprep kit (Zymo Research) according to the manufacturer’s protocol. NGS was performed on the Illumina MiSeq in the UAB Genomics Core Facility as described elsewhere (24). To identify mutational hot spots, the raw sequence fastq files were aligned to the reference genome using BWA version 0.7.17-r1188. Aligned reads were then sorted with Picard version 2.9.2 SortSam. Naïve Variant Caller (Galaxy version 0.0.4) called the variants and VariantAnnotator (Galaxy version 1.3.2) counted the bases at each position.

### RT-qPCR

Total RNA was extracted from 0.1 ml viral sample using Direct-zol™ RNA Miniprep kit (Zymo Research) according to the manufacturer’s protocol. cDNA was synthesized using random primers and QuantiTect Reverse Transcription kit (Qiagen). Quantitative PCR (qPCR) reactions were performed in a final volume of 20 µl containing *SsoFast™ EvaGreen*® *Supermix* (Bio-Rad) using CFX96 Real-Time PCR Detection System (Bio-Rad) and 500 nM SARS-Cov-2 nsp1-specific CCTCAACTTGAACAGCCCTATG forward and GAATGCCTTCGAGTTCTGCTAC reverse primers.

### Molecular Modeling

Models for the complete SARS-CoV-2 S ectodomain trimer and trimer with the GLTSKRN insertions after amino acid residue 214 were generated with SWISS-MODEL (25) based on the coordinates of PDB ID: 7JJI (26). Additional models of S trimers individually harboring I68R, N74K or S247R mutations were generated using COOT (27). The structures were visualized in BIOVIA Discovery Studio Visualizer.

### Biosafety

Virus rescue and analyses, which required infectious virus, were performed in BSL3 SEBLAB facility of the University of Alabama at Birmingham according to the protocols approved by the Institutional Biosafety Committee.

## RESULTS

The original WA1/2020 isolate that we received from WRCEVA (UTMB) had already been passaged 4 times (P4). Similar to another previously described P4 stocks (28, 29), it produced heterogeneous plaques on Vero E6 cells (Fig. 1A), suggesting presence of multiple variants. In addition, plaques formed on Vero E6 cells were not clear, because some cells in the plaque areas remained uninfected and continued to grow. Cloning of the cells showed that Vero E6 were heterogeneous in terms of ACE2 expression. Fig. 1B presents an example: clone 1 of Vero E6 demonstrated a 15-fold higher level of ACE2 expression than clone 2 derived from the same cell line. To avoid a time-consuming selection of clones with the highest level of ACE2, we generated a cell line of Vero E6 cells, which stably expressed human ACE2 (hACE2) in addition to the endogenously expressed homolog (see Materials and Methods for details). The increase in ACE2 expression in Vero/ACE2 cells (Fig. 1A) resulted in formation of noticeably larger and clear plaques, compared to those developed on the parental Vero E6 cells. One of the clones of Vero/ACE2 cells (cl. 7) (Fig. 1B) was used in most plaque assays. Similar ACE2-expressing stable cell lines were also generated on human Huh7.5, hamster BHK-21 and mouse NIH 3T3 cells. All these cell lines became highly susceptible to SARS-CoV-2 and, upon infection, rapidly developed a cytopathic effect (CPE). However, in contrast to Vero/ACE2, infection of these cells with WA1/2020 induced formation of very large syncytia with hundreds of cells involved. This resulted in relatively low virus production, and thus, limited application of the latter cell lines.

**FIG. 1.**
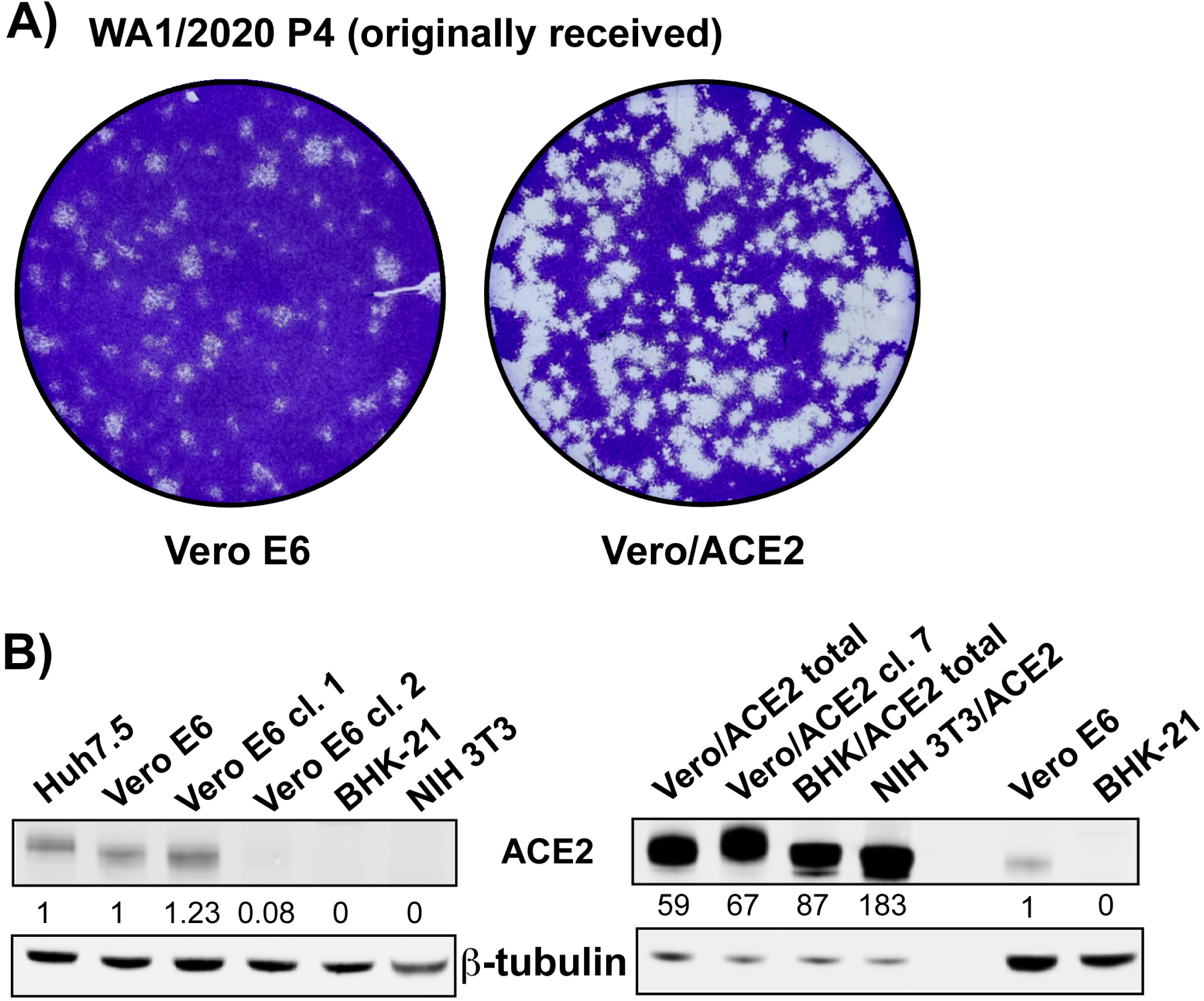
The original sample of WA1/2020 is heterogeneous. (A) The original WA1/2020 (P4) isolate received was titrated in parallel on Vero E6 and Vero/ACE2 cells (see Materials and Methods for details). Cells were fixed and stained at 68 h p.i. (B) Expression of ACE2 in different cell lines and cell clones. “Total” indicates that cells were not cloned after blasticidin selection.

Attempts to isolate variants from the pinpoint and small plaques (Fig. 1A) developed by the WA1/2020 P4 virus on Vero/ACE2 cells were unsuccessful. As it had been previously shown (29), plaque-purified variants immediately evolved during amplification in both Vero E6 and Vero/ACE2 cells, and harvested stocks produced heterogeneous plaques. This was an indication of rapid evolution of SARS-CoV-2 during its passaging on Vero cells.

In order to better understand the mechanism of SARS-CoV-2 evolution in cultured cells, virus from the original stock was passaged 5 times on Vero E6 with gradual decrease of the inoculum volume used for infection. Infectious titers of the harvested stocks increased from 1.5×10^6^ PFU/ml at starting passage 5 to 1.5-2×10^8^ PFU/ml at passage 9. After this selection, the harvested viral pool became homogeneous in terms of plaque size (Fig. 2A), and, compared to the original stock, the plaques formed became dramatically larger (Figs. 1A and 2A). Comparison of particle concentrations released into the media [genome equivalents per ml (GE/ml)] demonstrated that passaging did not result in a profound increase in virus release, because concentrations of GE in the media at different passages were similar (Fig. 2B). This passaging drastically decreased the GE:PFU ratio, suggesting that the ∼100-fold higher infectious titers resulted from a higher infectivity of the released virions. However, it remained unclear whether the more infectious variant(s) was present in the originally received, heterogeneous pool of SARS-CoV-2, or there was viral evolution and selection during the above-described passaging.

**FIG. 2.**
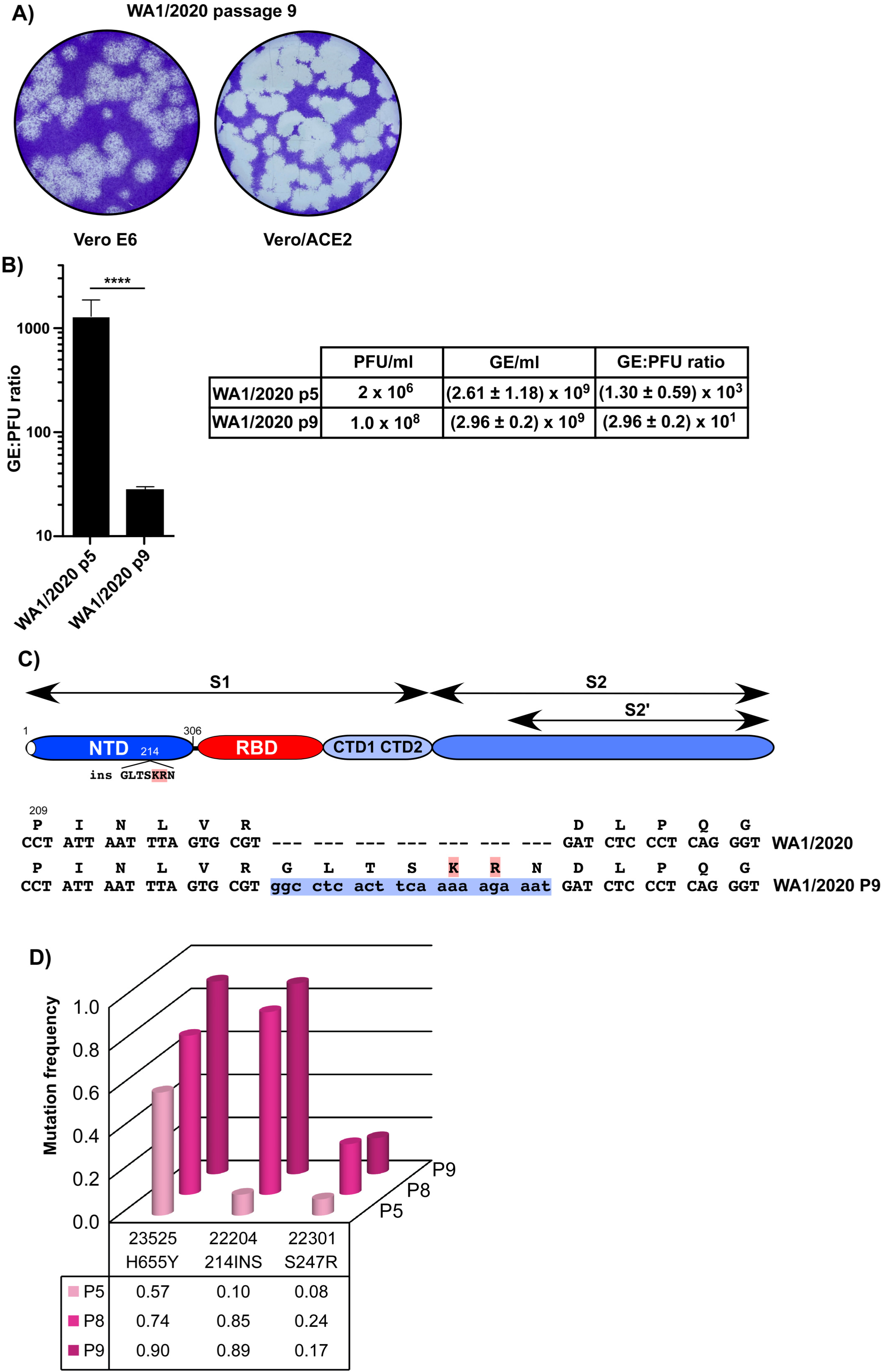
WA1/2020 isolate evolves to the large plaque phenotype and higher infectivity (A) Plaques formed by WA1/2020 virus on Vero E6 and Vero/ACE2 cells. Plaques were stained by crystal violet at 68 h p.i. (B) Concentrations of GE and PFU, and GE:PFU ratios in the samples of WA1/2020 at P5 (passage 1 following receipt) and P9 (passage 5 following receipt). Significance of differences was determined by the unpaired t-test with Welch’s correction (**\*\*\*\***, *P***<**0.0001; n**=**3). (C) The schematic presentation of the SARS-CoV-2 spike protein, position of the identified insertion, and its nt and aa sequences. (D) Evolution of WA1/2020 pool during virus replication in Vero E6 cells for 5 passages.

### Evolution of the S protein in WA1/2020 variant

Next, the S protein-coding sequence of the P9 viral pool was amplified by RT-PCR and sequenced. The furin cleavage site (FCS), which was frequently deleted in prior studies (29), was found to be intact, indicating that deletion of this sequence is not the only means of increasing infectious titers *in vitro*. A major mutation identified in the S protein of passage 9 virus was an in-frame insertion of 21 nt into the NTD-coding sequence (Fig. 2C). This insertion added a peptide GLTSKRN between aa 214 and 215. A second dominant mutation, H655Y, was in the CTD2 domain. It has been found in many natural isolates and became dominant in Brazilian lineage P.1 (https://www.gisaid.org/hcov19-variants).

Further analysis of viral pools, which were harvested at different passages, by Next Generation Sequencing (NGS) demonstrated that the insertion was likely present in the originally received WA1/2020 P4 sample and rapidly became dominant in the viral pool by passage 9 during further passaging (Fig. 2D). Thus, the insertion-containing variant was selected from the original heterogeneous pool, as it had higher infectivity for Vero E6 cells and, thus was able to spread more efficiently than others. The distinguishing characteristic of the inserted peptide was the presence of 2 positively charged aa, lysine and arginine. Their presence could potentially increase affinity of the virus to GAG(s), which are abundantly present at the plasma membrane, and mediate more efficient primary viral attachment to the cells before interaction with the ACE2 receptor. The H655Y mutation was already abundant in P5 virus, and thus, could not directly increase viral infectivity, at least while being alone. In addition, we have also detected an S247R substitution in all the samples of passaged WA1/2020 (Fig. 2D). It was less abundant than the above insertion and H655Y mutation, present in smaller fraction of viral genomes, and its frequency was not increasing. However, the possible benefit of S247R for viral infectivity and spread cannot be completely ruled out.

### cDNA-derived SARS-CoV-2 rapidly evolves

In another line of experiments, we analyzed evolution of SARS-CoV-2 derived from recombinant cDNA. This cDNA was synthesized according to the GenBank sequence NC_045512.2, but the ORF 7a was replaced with a GFP-coding sequence (Fig. 3A). GFP expression was used to simplify monitoring of infection spread. SARS-CoV-2/GFP (CoV-2/GFP) (Fig. 3A) was propagated in two low copy number plasmids (see Materials and Methods for details). Rescuing of the virus was performed by electroporation of the *in vitro*-synthesized RNA into BHK-21 cells stably expressing N protein of SARS-CoV-2 from the persistently replicating alphavirus replicon (see Materials and Methods for details). The transfected cells were seeded on a subconfluent monolayer of Vero/ACE2 cells and the recombinant virus was harvested after development of cytopathic effect (CPE). The initially harvested CoV-2/GFP expressed high levels of GFP (Fig. 3B) upon infection of Vero cells and, as in the case of WA1/2020 P4, formed heterogeneous small and medium size plaques (Fig. 3C). Its further passaging led to rapid evolution to the large plaque-forming phenotype (Fig. 3C). By passage 5, viral infectious titers increased by two orders of magnitude and approached 2×10^8^ PFU/ml. As with the above experiments using WA1/2020, the detected evolution did not result in more efficient accumulation of released particles, but rather increased their infectivity, which led to a strong decrease in the GE:PFU ratio (Fig. 3D).

**FIG. 3.**
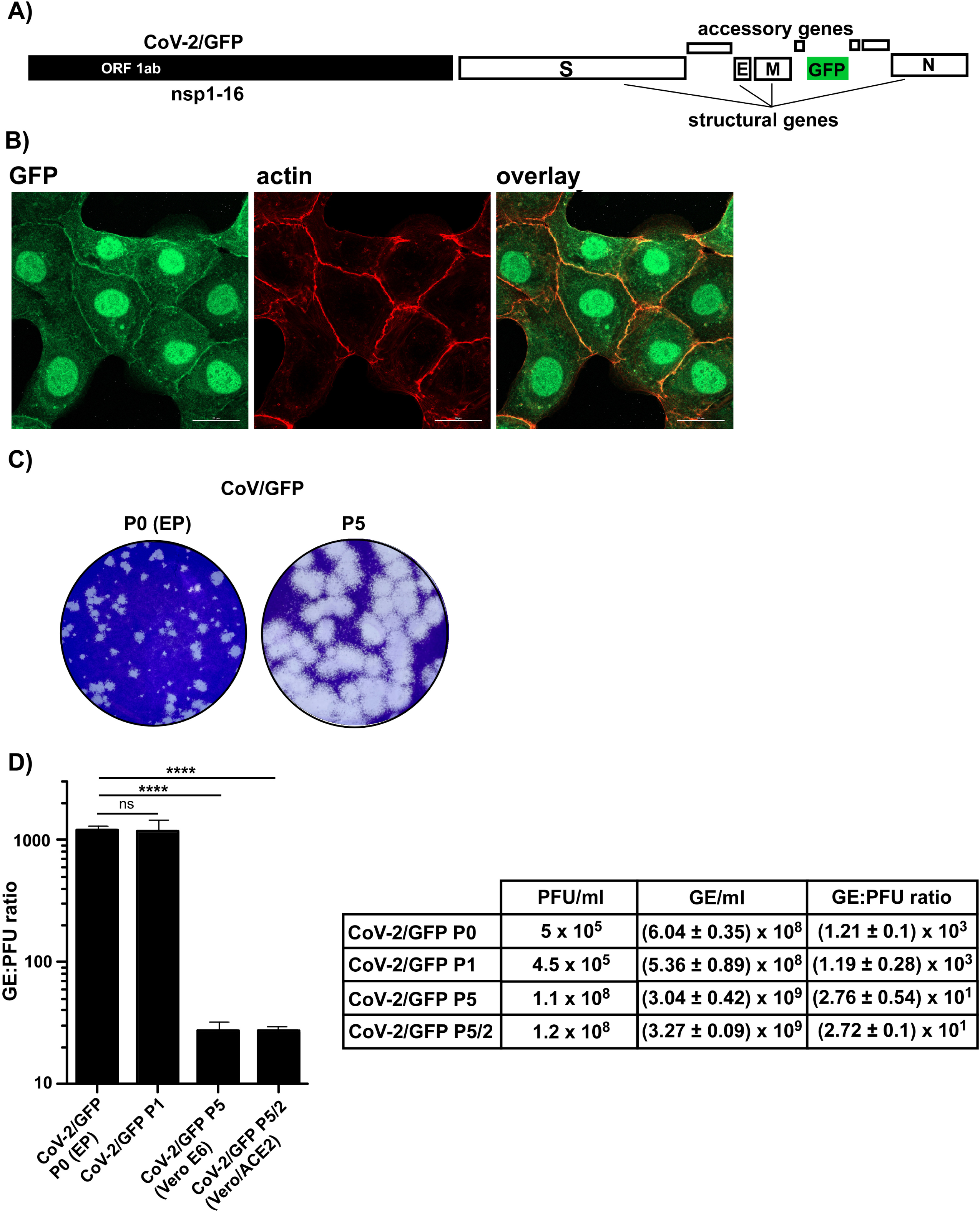
The cDNA-derived CoV-2/GFP rapidly adapts during passaging in Vero cells to higher infectivity and large plaque phenotype. (A) Schematic presentation of the coding strategy of the recombinant CoV-2/GFP genome. (B) Vero/ACE2 cells were infected with CoV-2/GFP at an MOI of 10 PFU/cell and fixed at 8 h p.i. with PFA and stained with phalloidin, labeled with Alexa Fluor 546. Images were acquired on a Zeiss LSM 800 confocal microscope in Airyscan mode with a 63X 1.4NA PlanApochromat oil objective. (C) Plaques formed on Vero/ACE2 by CoV-2/GFP P0 (harvested after electroporation) and by the same virus after 5 passages on Vero/ACE2 cells. Staining was performed at the same time p.i. (D) Concentrations of GE and PFU, and GE:PFU ratios in the samples of CoV-2/GFP harvested at P0, P1 and P5 on Vero E6 cells. P5/2 indicates a sample harvested at passage 5 on Vero/ACE2 cells. Significance of differences was determined by the unpaired t-test with Welch’s correction (**\*\*\*\***, *P***<**0.0001; n**=**3).

A preliminary Sanger sequencing of the S gene in the high passage viral pool identified a single nucleotide mutation resulting in S686G substitution, which was the first aa downstream of the FCS (Fig. 4A). Further NGS-based analysis demonstrated rapid accumulation of the variants with the latter mutation in viral samples harvested at different passages (Fig. 4B). More than 60% of the variants in the P1 pool already contained this mutation. The P7 pool had the latter mutation in nearly 100% of the genomes. This suggested that the S686G substitution had a stimulatory effect on viral spread in Vero cells. There was also a gradual increase in the presence of additional mutations, mostly in the NTD of the S protein. Two of them, S68R and N74K, increased the positive charge of the surface of this domain. Another mutation in the furin cleavage cite (R685H) was detectable at early passages and likely inactivated the furin-mediated cleavage. However, it was eliminated by passage 7, suggesting that the latter mutation was less advantageous than S686G for viral spread. In the course of this study, electroporation of the *in vitro*-synthesized SARS-CoV-2/GFP RNA was repeated multiple times, and there was always evolution of the rescued virus to a large plaque-forming phenotype within very few passages. In addition, we have also sequenced genomes in the virions pooled together from three independent electroporations of the *in vitro*-synthesized CoV-2/GFP RNAs and found that rescued viruses also contained mutations in FCS site (S686G at 59% and R683P at 30%). Thus, as in the case of the above described natural isolate WA1/2020, serial passaging of the cDNA-derived virus *in vitro* was also leading to rapid selection of variants demonstrating higher infectivity and formation of large plaques. Albeit this adaptation was achieved by accumulation of different mutations in the spike-coding gene. In the case of recombinant virus, the mutations in FCS became dominant by passage 2 and were likely selected from those randomly generated by the SP6 polymerase during RNA synthesis.

**FIG. 4.**
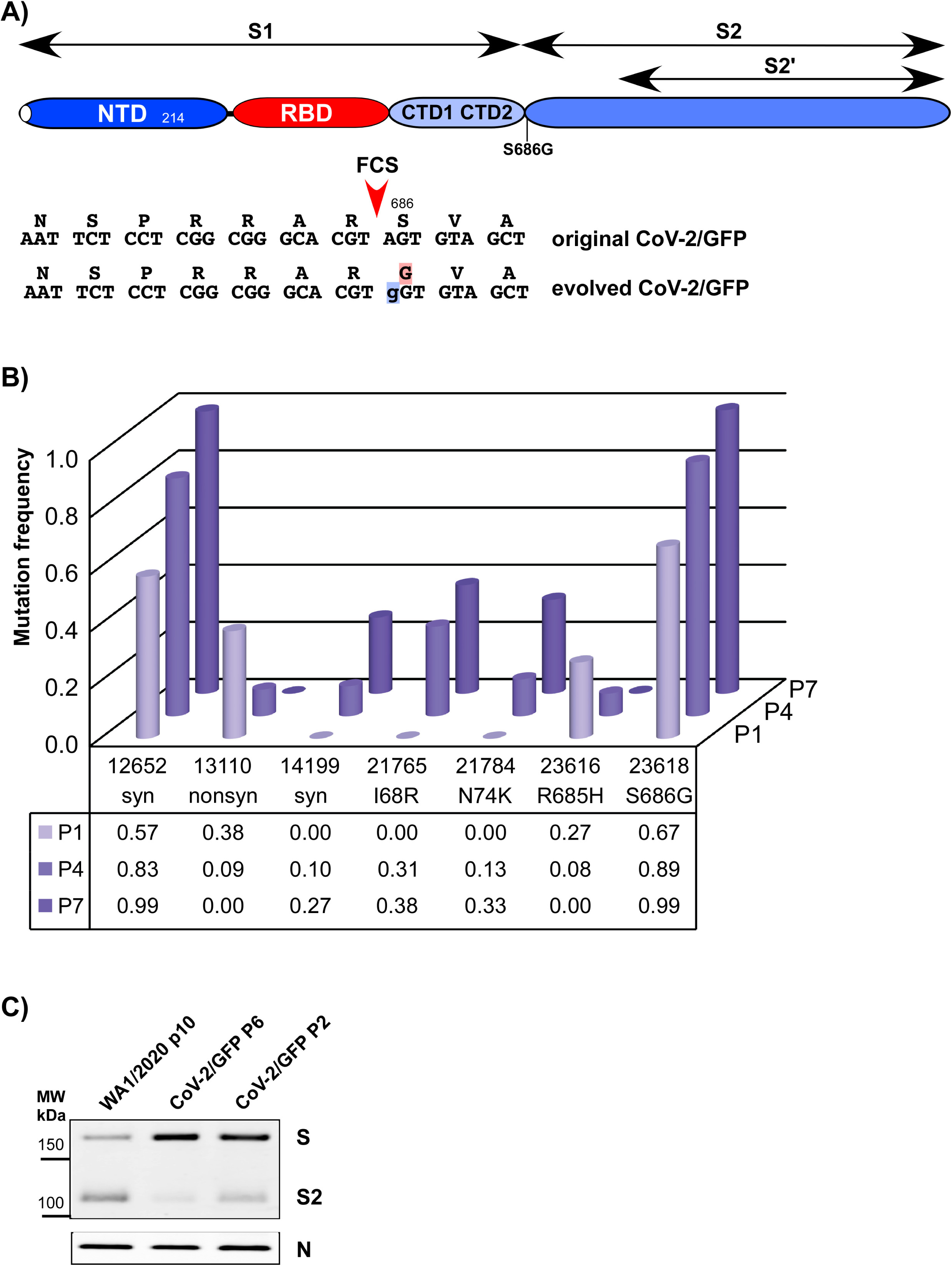
CoV-2/GFP acquires an S686G mutation in P1’ of FCS. (A) The schematic presentation of the SARS-CoV-2 S protein, position of the mutation, and nt and aa sequences of the gene fragment in the parental and Vero cells-adapted variant. (B) Evolution of CoV-2/GFP pool during virus replication in Vero E6 cells for 5 passages. (C) Western blot analysis of S protein in the particles released from cells infected with WA1/2020 and CoV-2/GFP harvested at different passages on Vero E6 cells (see Materials and Methods for details).

Next, we experimentally evaluated the effect of the S686G mutation on processing of the S protein in released viral particles. The viruses analyzed included i) WA1/2020 P10, ii) rescued CoV-2/GFP P2 and iii) P6 of the latter virus. All the samples were produced in serum-free media (see Materials and Methods for details), and the released particles were harvested at 24 h p.i., before CPE development. Samples of the media were used directly for Western blotting. The results presented in Fig. 4C demonstrated that the insertion in the NTD and H655Y mutation identified in high passage WA1/2020 did not abrogate furin-mediated processing of S protein. However, the S686G mutation, which is located in the P1‘ position of the FCS, strongly affected the efficiency of cleavage, and most of S protein in the released virions remained unprocessed.

### Mutations identified in the high passage viruses strongly increase infectivity of the cDNA-derived recombinant CoV-2/GFP

The above data suggested that serial passaging of SARS-CoV-2 *in vitro* leads to evolution of its spike protein that results in higher infectious titers. To experimentally support these data, we introduced the most prevalent mutations identified into cDNA of CoV-2/GFP. One of the variants encoded the above-described insertion in its NTD (CoV-2/GFP/ins), the second one had an S686G mutation downstream in P1’ position of FCS (CoV-2/GFP/G), and the third mutant (CoV-2/GFP/ins/G) contained both modifications in the S protein. Positive effects of the mutations on viral spread were already detectable during virus rescue from the *in vitro*-synthesized RNAs. After electroporation, cells that received mutant RNAs developed profound CPE within 48 h, while the parental construct SARS-CoV-2/GFP produced CPE after 72 h. Sequencing of the S gene in the rescued viruses confirmed the presence of introduced mutations in the genomes. The recombinant viruses formed large plaques on Vero/ACE2 cells (Fig. 5A), which were similar to those of the parental, serially passaged viruses (Figs. 2A and 3C). Their larger size correlated with the strong decrease in GE:PFU ratios suggesting their higher infectivity for Vero cells (Fig. 5B). Importantly, the combination of both mutations further reduced the GE:PFU ratio. Comparative analysis of viral replication rates in Vero/ACE2 cells (Fig. 5C) showed that at any times p.i., the mutations led to higher infectious titers identical to those of the originally selected, late passage viruses. The recombinant viruses, but not CoV-2/GFP P0 and WA1/2020 P4, also caused complete CPE by 40 h p.i. The double mutant CoV-2/GFP/ins/G replicated more efficiently than the single mutants (Fig. 5C). Mutations in the S proteins of WA1/2020 P9, CoV-2/GFP P5 and CoV-2/GFP/ins/G P0 did not abrogate infectivities of the viruses for Calu3 cells (Fig. 5C), which are widely used in SARS-CoV-2 research. Variants containing the S686G mutation (the late passage CoV-2/GFP, CoV-2/GFP/G and CoV-2/GFP/ins/G) demonstrated less efficient processing of the S protein (Fig. 5D). Taken together, the data indicated independent functions of the mutations introduced into S protein in viral replication to higher infectious titers, at least in Vero cells. However, it remained unclear whether these changes in the S protein function via the same or different mechanisms.

**FIG. 5.**
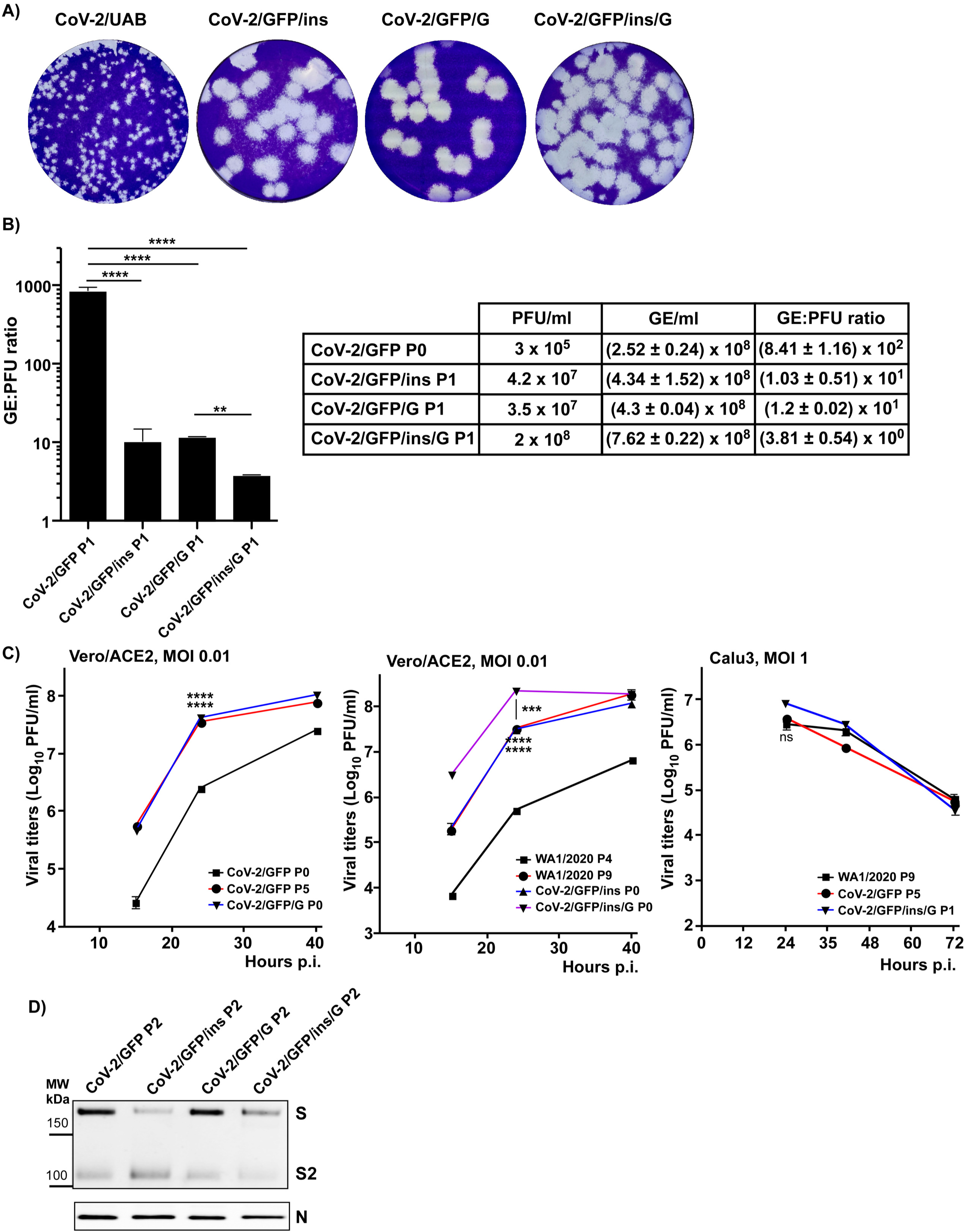
Mutations in the S protein-coding sequence confirm the large plaque phenotype and higher infectivities of the recombinant viruses to Vero cells. (A) Plaques formed by the early passage (P1) isolate of CoV-2/UAB and the recombinant CoV-2/GFP-based variants, containing either the insertion in the NTD or S686G mutation, or both modifications. In all cases, P1 viruses were assessed in the standard plaque assay and plaques were stained at day 3 p.i. (B) Concentrations of GE and PFU, and GE:PFU ratios in the samples of CoV-2/GFP encoding the S protein with the indicated modifications. (C) Vero/ACE2 cells were infected with the indicated viruses at an MOI of 0.01 PFU/cell. Calu3 cells were infected at an MOI of 1 PFU/cell. After 1 h of infection, cells were washed 3 times with PBS and further incubated in the corresponding complete media. At the indicated time points, media were replaced, and viral titers were determined by plaque assay on Vero/ACE2 cells. For B and C significance of differences was determined by the unpaired t-test with Welch’s correction (**, P<0.01; ***, P<0.001; **\*\*\*\***, *P***<**0.0001; n**=**3). (D) Western blot analysis of S protein in the particles released from Vero cells infected with CoV-2/GFP, CoV-2/GFP/ins, CoV-2/GFP/G and CoV-2/GFP/ins/G (see Materials and Methods for details).

### Mutations in the S protein of selected mutants enhance SARS-CoV-2 binding to heparin

Next, we evaluated interaction of different viral variants with heparin. Samples of all of the viruses indicated in Fig. 6 were prepared in serum-free media. This set included the UAB clinical isolate CoV-2/UAB P2, WA1/2020 P5, early passage CoV-2/GFP P2, late passage WA1/2020 P10 and CoV-2/GFP P7. Viral samples were diluted to 0.1 M concentration of NaCl, loaded to the heparin sepharose column and eluted using a step gradient of NaCl (see Materials and Methods for details). The results presented in Fig. 6 demonstrate that the low passage samples of natural isolates WA1/2020 P5 and CoV-2/UAB had low affinities to heparin and mostly remained in the flow through (FT) fraction containing 0.1 M NaCl. Smaller peaks of infectious viruses were also eluted by 0.2 M NaCl, but not in other fractions with higher salt concentrations. CoV-2/GFP P2 demonstrated noticeable heterogeneity in terms of binding to heparin. Some of the virus was still present in the FT fraction and the rest was eluted with 0.25-0.3 M NaCl. Passaged viruses WA1/2020 P10 and CoV-2/GFP P7 bound to heparin sepharose more efficiently, and less than 1 % of the infectious viruses were found in the FT fraction. WA1/2020 P10 and CoV-2/GFP P7 were eluted by 0.35 M and 0.3 M NaCl, respectively. Thus, the mutations accumulated in the S protein of the high passage viruses, which became more infectious for Vero cells, also increased their affinities to heparin.

**FIG. 6.**
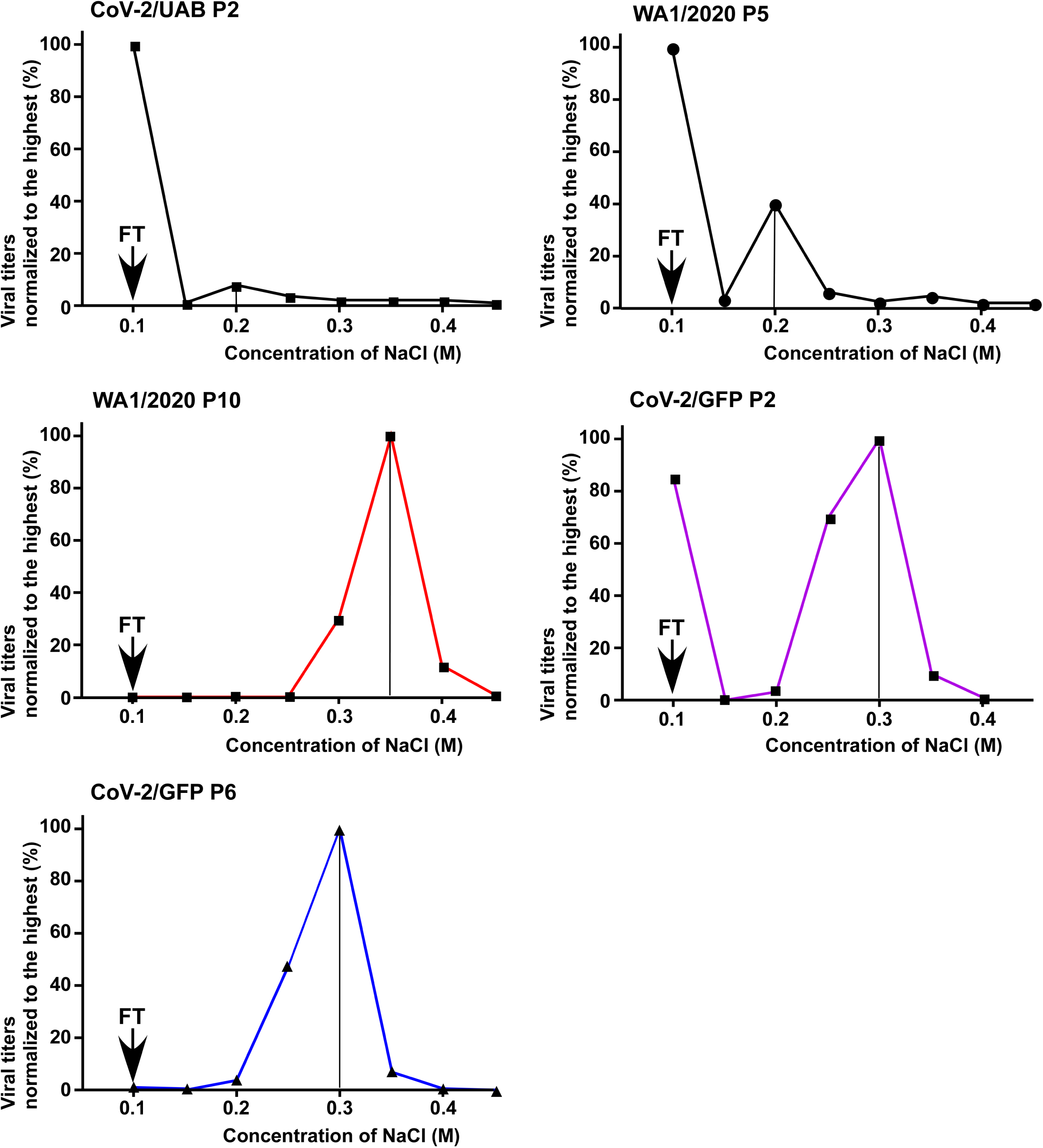
After passaging on Vero cells, SARS-CoV-2 acquires higher affinity to heparin sepharose. Samples of the indicated viruses were prepared in the serum-free media, diluted to 0.1 M NaCl and loaded to a column of heparin sepharose. After washing with 0.66 x PBS, which contains 0.1 M NaCl, viruses were eluted with PBS, containing increasing concentrations of NaCl. Viral titers in each fraction, including FT, were determined by plaque assay on Vero/ACE2 cells. Titers were normalized to those in the fraction with the highest viral concentration. The experiments were repeated twice with reproducible results. The data from one of the experiments are presented.

The pools of the high passage viruses still contained variants with a variety of mutations in the S protein-coding sequence. Therefore, it was difficult to make definitive conclusions about the effects of GLTSKRN insertion and S686G mutation on virions’ affinity to heparin. Therefore, we also evaluated binding of the above-described designed recombinant viruses to heparin sepharose. Based on sequencing data, they had no additional aa substitutions in the S proteins. Their samples were also prepared in serum-free media. Binding and elution patterns of CoV-2/GFP/ins P1 and CoC-2/GFP/G P1 (Fig. 7) correlated with those of the late passage WA1/2020 and CoV-2/GFP (Fig. 6), respectively. In the FT fraction, both recombinant viruses were present at very low levels. The peak of CoV-2/GFP/ins variant elution was at 0.35 M NaCl, and CoV-2/GFP/G was eluted by 0.3M NaCl. Thus, both the insertion and S686G mutation increased affinities of the viruses to heparin sepharose. The S686G mutation did not increase the positive charge of the S protein surface, but it likely transformed the unprocessed furin cleavage site into the heparin-binding site. Interestingly, the double mutant CoV-2/GFP/ins/G, which contained both the insertion in the NTD and S686G mutation, was also eluted by 0.35 M NaCl. Thus, the combination of both modifications in the spikes did not increase viral affinity above that of the high-passage natural isolates. This was suggestive that although binding to heparin was likely determined by two separate sites, the higher affinity was mostly determined by the insertion in the NTD and played the dominant role.

**FIG. 7.**
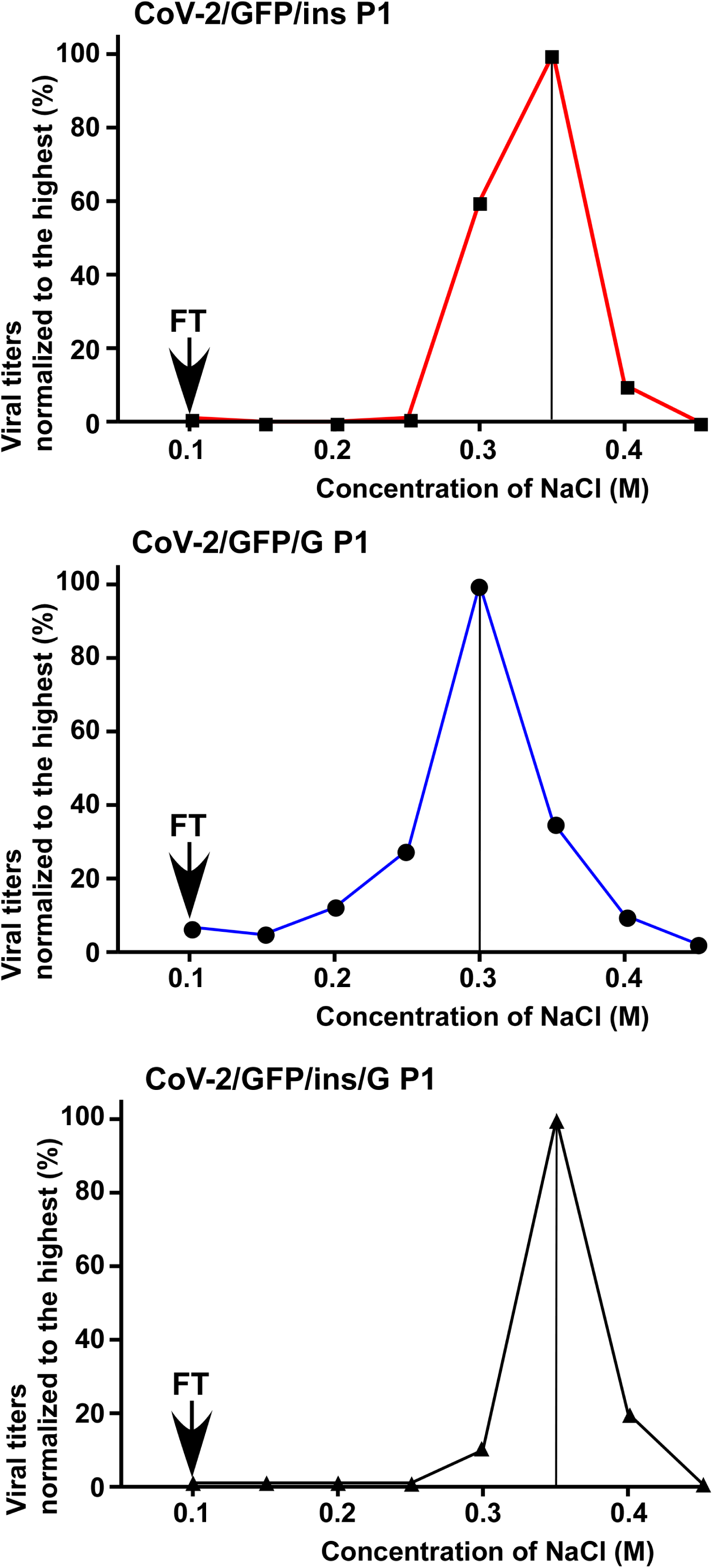
Recombinant viruses with the indicated mutations in the S protein exhibit high affinity to heparin sepharose. Viral samples were prepared in serum-free media and analyzed as described in the legend to Fig. 6. Viral titers in each fraction, including FT, were determined by plaque assay on Vero/ACE2 cells. Titers were normalized to those in the fraction with the highest concentration of infectious virus. The experiments were repeated twice with reproducible results. Data from one of the representative experiments are presented.

Taken together, the results of these experiments demonstrated that passaging of SARS-CoV-2 leads to its rapid evolution towards a more efficient binding to heparin. These adaptive mutations include insertion of the short positively charged peptide after aa 214 in the S protein and transformation of the furin cleavage cite into a heparin-binding aa sequence. However, additional contributions of other mutations in the NTD of the spike protein, such as S68R or N74K, which may additionally increase its affinity to HS, are also possible.

Variants having the S686G mutation exhibited one more interesting characteristic. They lost the ability to form syncytia on Vero/ACE2 cells (Fig. 8). In our experimental conditions, SARS/UAB, WA1/2020 P9 and CoV-2/GFP/ins formed very large syncytia. The early passage CoV-2/GFP P2, which had a mixed population of viral variants, was still capable of forming at least small syncytia, but by passage 7, it completely lost this ability. Both CoV-2/GFP/G and double mutant CoV-2/GFP/ins/G were unable to produce syncytia at all.

**FIG. 8.**
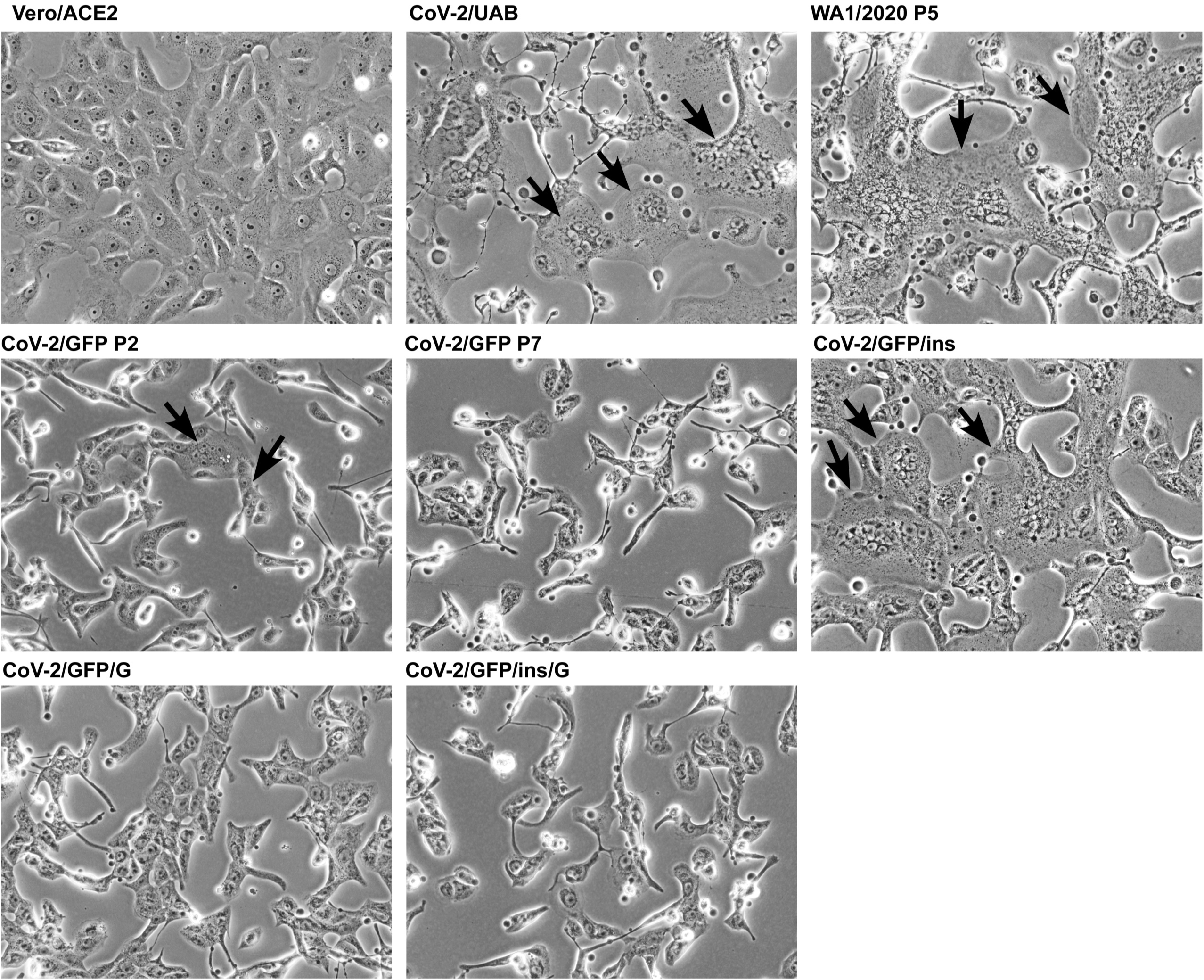
The S686G mutation has negative effect on the ability of SARS-CoV-2 virus to form syncytia. Vero/ACE2 cells in 6-well Costar plates were infected with the indicated variants at an MOI of 2 PFU/cell and incubated in serum-free media for 16 h at 37°C. Then they were fixed by 4% PFA, and images were acquired on EVOS microscope. Syncytia are indicated by black arrows.

## DISCUSSION

In this report, we demonstrate that SARS-CoV-2 rapidly adapts to replication in Vero E6 cells, which are most commonly used for virus propagation. This adaptation results in higher infectious titers *in vitro*, which are determined by higher infectivity of the viral particles rather than by more efficient particle release. The evolved variants form dramatically larger plaques on Vero cells and demonstrate more rapid infection spread. Similar evolution of SARS-CoV-2 to the large plaque-forming phenotype has been detected in previous publications, suggesting that this is a common effect, and likely independent of experimental conditions (28).

Viral evolution developed by two mechanisms. The first means of adaptation was determined by the insertion of a 7-aa-long peptide into the S protein of the WA1/2020 isolate (Fig. 2). It was already present in the provided heterogeneous viral sample, and rapidly became dominant during further passaging. This GLTSKRN insertion was located in the NTD (Figs. 2, 9 and 10), suggesting an important role of the NTD in primary attachment of the virus to cells prior to interaction with the ACE2 receptor. The inserted peptide contained two positively charged aa and made both the evolved natural variant and its cDNA-derived analog more infectious *in vitro*. It strongly increased affinity of both viruses to heparin sepharose (Figs. 6 and 7) and decreased their GE:PFU ratios by 2 orders of magnitude compared to those of the low passage natural isolates. Consequently, it had also positive effects on the plaque size and rates of infection spread. The identified insertion increased positive charge on the surface of the NTD domain (Figs. 9 and 10), and this explained the higher affinity of the mutant virus to heparin. A small fraction of passaged WA1/2020 contained another mutation, S247R, which also contribute to an increase in the positive charge of the NTD surface (Fig. 9). Since presence of the latter mutation did not increase during passaging, its contribution to infection spread was likely lower than that of the above-described insertion.

**FIG. 9.**
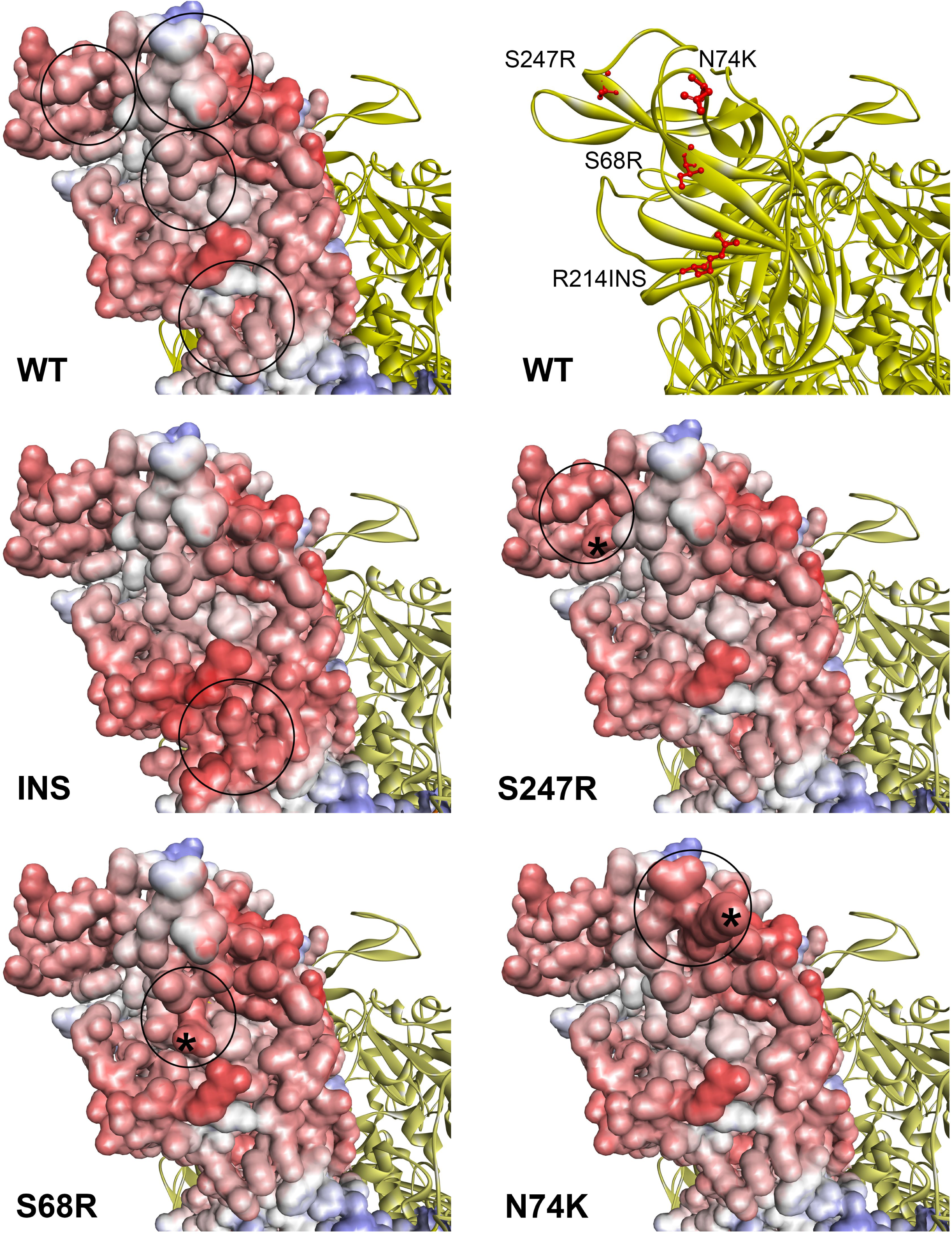
Mutations acquired by the NTD during virus passaging in Vero cells increase the positive charge of the protein surface. Electrostatic surface rendering is presented. Blue and red indicate electronegative and electropositive surfaces, respectively. Positions of the indicated aa substitutions and peptide insertion GLTSKRN between amino acid residues 214 and 215 are shown by black circles and asterisks. Images were generated with BIOVIA Discovery Studio Visualizer.

**FIG. 10.**
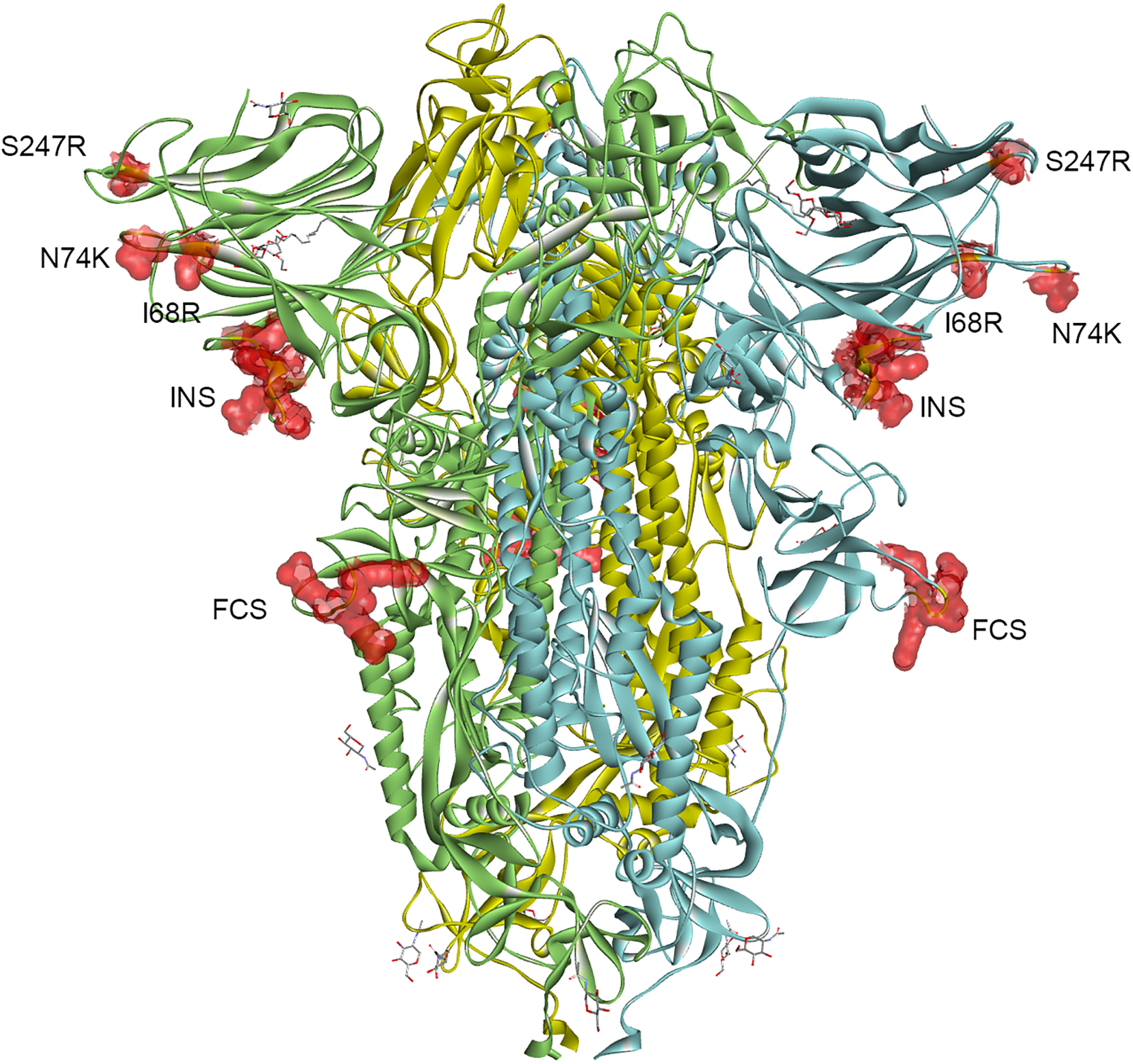
The 3D model of the trimeric SARS-CoV-2 spike ectodomain is shown. The individual spike subunits are shown in green, yellow and blue colors. Locations of the single mutations, the insertion in the NTD, and uncleaved FCS are highlighted with red spheres.

The second mechanism of SARS-CoV-2 adaptation was based on the appearance of the S686G substitution in P1’ position of the S1/S2 cleavage site, FCS, which was rapidly selected in the S protein of recombinant CoV-2/GFP. The previously published data showed that the peptide representing FCS is often entirely or partially deleted during virus passaging (28–30), and the same S686G mutation was also found after passaging of other SARS-CoV-2 isolates (28, 29). The identified mutation strongly affected processing of the S protein (Figs. 4C and 5D) and formation of the syncytia (Fig. 8) during viral replication. It also increased infectivity of harvested viruses, their plaque sizes, and infectious titers (Figs. 3C and 5C). In prior studies, the FCS could be inactivated by either its deletion or accumulation of point mutations eliminating the arginines essential for furin cleavage (29). Viral mutants with deleted FCS, and R682L, S686G and H655L mutations have been detected in the stock of WA1/2020 isolate from CDC (31). The effect of the FCS deletion was experimentally evaluated in the context of pseudovirions or recombinant SARS-CoV-2 (32, 33). It resulted in higher infectious titers of the viruses harvested from Vero cells, but the detected increase was small. It has also been demonstrated that the deletion of FCS affected viral infectivity in human cells including, Calu-3, Caco2 and primary lung epithelia cells.

In our study, the selected recombinant viruses with the S686G mutation replicated to two orders of magnitude higher infectious titers in Vero cells than did the natural isolates and previously designed deletion mutants. They also remained capable of infecting and reaching high titers on Calu-3 cells (Fig. 5). These results suggest that in contrast to FCS deletion, inhibition of the S1/S2 processing by S686G is likely not the only advantage of the new variants. Viral spikes with S686G substitutions retained the entire polybasic motif, which was originally required for cleavage by furin protease (Fig. 4A), and such viruses attained high affinity to heparin sepharose (Figs. 6 and 7), higher infectivity and more efficient spread on Vero cells. Similar evolution of the FCS into the heparin-binding motif has previously been described for some of the group 1 coronaviruses (34). In that study, the resultant polybasic motif of the S protein-specific FCS was functioning in HS-binding. Interestingly, mutations in FCS have been shown to both increase and decrease CoV pathogenicity. HCoV-OC43 encodes the RRSRG cleavage site, which is similar to that in our S686G mutant, and its spike protein remains unprocessed (35). However, the clinical isolates with functional FCS were viable and showed lower neurovirulence (36). On the other hand, in the feline enteric coronavirus (FECV), FCS mutations abolishing spike protein processing transformed the virus to another more pathogenic biotype, feline infectious peritonitis virus (FIPV) (37, 38). In our study, at the early passages, we were able to detect another CoV-2/GFP variant with mutated positively charged aa in the furin cleavage site (R685H mutation). However, it was likely less competitive than S686G mutant, and was eliminated from the viral pool within a few subsequent passages. Thus, the S686G mutant apparently produced higher infectious titers than other variants with mutated arginine-rich motifs. Interestingly, the latter RRAR remains in the S1 subunit of the original isolates after furin-mediated processing. However, in this post cleavage conformation, it likely interacts with heparin less efficiency, if at all, than that in the uncleaved S protein of the selected mutant. Interestingly, downregulation of the S protein processing had a negative effect on the ability of the recombinant viruses to induce syncytia formation at least in Vero cells (Fig. 8), but it remains unclear whether the mutants became less efficient in forming syncytia *in vivo*.

The experimental evidences suggest that the virus with unprocessed FCS remains less efficient in HS binding than the variant with the NTD insertion. First, viruses with S686G are eluted from heparin sepharose by lower concentrations of NaCl (Figs. 6 and 7). Secondly, in contrast to the insertion mutant, the originally selected CoV-2/GFP variant with the S686G mutation continued to evolve and attain additional mutations in the NTD, such as I68R and N74K (Figs. 4 and 10). Their accumulation was relatively slow, but clearly detectable, and implied their additional contribution to HS binding and viral infectivity. Analysis of sequences submitted to GISAR revealed that accumulation of mutations that increase the positive charge of NTD upon passaging in Vero cells, is quite common: N74K was detected in EPI_ISL_1039208, EPI_ISL_1190402, EPI_ISL_1718321, EPI_ISL_1707039, EPI_ISL_1785073, EPI_ISL_1785076, and I68R/K was found in EPI_ISL_493139, EPI_ISL_2226226. The WA1/2020 from CDC was reported to contain 9 low frequency R/K substitutions in NTD including a 4-aa-long insertion after D215 (31). The indicated I68R and N74K mutations, which additionally increase the positive charge of the NTD surface (Fig. 9), are also located relatively close to the above-described S247R (Fig. 10). Currently, there is not enough information to speculate whether the increase of positive charge on the NTD surface attenuates the virus, makes it more pathogenic or has no effect on pathogenicity. Importantly, since R/K mutations in NTD are highly beneficial for virus propagation *in vitro*, viruses used for animal studies should be thoroughly analyzed by NGS.

Combining the positively charged insertion and S686G mutation in the same S protein did not detectably increase the affinity of infectious viral particles to heparin sepharose above the level determined for CoV-2/GFP/ins (Fig. 7). However, the recombinant double mutant demonstrated significantly higher infection rates than did the single site mutants (Fig. 5). CoV-2/GFP/ins/G was reaching the peak titers above 10^8^ PFU/ml by 24 h p.i. even at low MOI, such as 0.01 PFU/cell. This was an indication that during viral infection, the introduced mutations, GLTSKRN insertion and S686G substitution, were able to function in either synergistic or additive modes. The structural model of the S protein supports the possibility of a combined effect. The above insertion and furin cleavage site are in close proximity on the surface of the S protein (Fig. 10) and may have a stimulatory effect on primary binding of the virus to GAGs at the plasma membrane. An important characteristic of the double mutant is that its further evolution in cultured cells appears to be unlikely. The latter virus demonstrated the lowest GE:PFU ratio, and its further decrease at least in Vero cells appears to be an improbable event.

Similar rapid evolution to more efficient spread *in vitro* was previously described for alphaviruses. As with SARS-CoV-2 in this study, chikungunya, Venezuelan equine encephalitis, Ross River and Sindbis viruses acquired 1 or 2 additional basic aa in the E2 glycoprotein, which had strong positive effects on virus interaction with HS (12-14, 39, 40). The mutations increased plaque sizes, stimulated spread of infection and dramatically decreased the GE:PFU ratio of the evolved alphaviruses. These mutant viruses were usually less pathogenic in mice. Therefore, it is tempting to speculate that, similar to other RNA+ viruses that adapted to binding to HS, the designed single mutants and the double mutant, in particular, became attenuated *in vivo*, but this hypothesis needs experimental support. Of note, the development of the recombinant live attenuated vaccine for SARS-CoV-2 will also require its large-scale production and passaging in cell culture, most likely in Vero cells. Therefore, viral evolution to a more stable HS-binding phenotype through the acquisition of mutations in the NTD is likely an unavoidable event and needs to be considered. The original isolates form very small plaques and have very high GE:PFU ratio. Their passaging results in rapid evolution of viral pool and selection of variants with higher infectivities *in vitro*. However, it is unlikely that they fully reproduce SARS-CoV-2 infection and pathogenesis *in vivo*.

Interestingly, previously published data, which were generated on the receptor-binding domain (RBD) and the isolated S protein, suggested that the RBD is capable of efficient interaction with heparin sepharose and HS (10). In our experiments, the early passage natural isolates demonstrated relatively inefficient binding to heparin sepharose. The main fraction of infectious virus remained in the FT fraction (0.1 M NaCl) and some virus was also eluted as a small peak at NaCl concentration of 0.2 M. However, the insertion- and/or S686G mutation-containing variants, which became adapted to cell culture, were eluted by NaCl at 0.35 and 0.3 concentrations, respectively, suggesting their higher affinities to heparin, which strongly correlated with viral infectivities and spread in cultured cells.

Importantly, we observed rapid adaptation of the recombinant, cDNA-derived virus, and the S686G mutation became dominant by passage 2, because it increased viral infectivity by 100-fold. The SP6 RNA polymerase used in our study and the T7 polymerase used by other groups have the mutation rates of 1-2×10^-4^/nt (41). Thus, the *in vitro*-synthesized viral genomes already contain a wide range of mutations, which represent a ground for further selection of the most efficiently spreading variants.

In conclusion, previously published data and the results of this study suggest that both, natural isolates of SARS-CoV-2 and the cDNA-derived recombinant variants, rapidly adapt to cell culture. They evolve to higher infectivity, which results in lower GE:PFU ratios, and more efficient infection spread by either acquiring mutations in the NTD, suggesting an important role of this domain in viral binding to the cells, or in the FCS. These mutations increase virus affinity to HS during its primary attachment to cells before interaction with the ACE2 receptor. The mutations leading to more efficient spread include i) substitutions of single aa by those positively charged, ii) insertions of short peptides that contain basic aa, and iii) S686G substitution that likely transforms FCS into the HS-interacting peptide. Such viral variants rapidly become dominant, and their further evolution and adaptations are unlikely. These adapted variants appear to be more convenient for CPE-based screening of the antiviral drugs. As with the HS-binding mutants of other RNA+ viruses, the evolved SARS-CoV-2 may also be attenuated *in vivo,* particularly the double mutant that demonstrates the most adapted phenotype and alterations in syncytia formation. Thus, they may also be used as a basis for development of stable live attenuated vaccines for COVID-19.

## ACKNOWLEDGMENTS

This study was supported by Public Health Service grants R01AI133159 and R01AI118867 to EIF, R21AI146969 to IF and U19AI142737 for TJG and by the UAB Research Acceleration Funds to EIF and IF.

